# Developmental exposure to a real-life environmental chemical mixture alters testicular transcription factor expression in neonatal and pre-pubertal rams, with morphological changes persisting into adulthood

**DOI:** 10.1101/2022.12.09.519746

**Authors:** Chris S. Elcombe, Ana Monteiro, Mohammad Ghasemzadeh-Hasankolaei, Vasantha Padmanabhan, Kevin D. Sinclair, Richard Lea, Neil P. Evans, Michelle Bellingham

**Affiliations:** School of Biodiversity, One Health, and Veterinary Medicine, College of Medical, Veterinary and Life Sciences, University of Glasgow, UK; Department of Pediatrics, University of Michigan, Ann Arbor, Michigan; University of Nottingham, Sutton Bonington Campus, Loughborough, UK

**Keywords:** Developmental toxicity, Reproductive toxicity, Environmental chemicals, Testicular dysgenesis syndrome, CREB

## Abstract

Environmental chemical (EC) exposure may be impacting male reproductive health. The translationally relevant biosolids treated pasture (BTP) sheep model was used to investigate gestational low-level EC mixture exposure on the testis of F1 male offspring. Adult rams from ewes exposed to BTP 1 month before and throughout pregnancy had more seminiferous tubules with degeneration and depletion of elongating spermatids, indicating “recovery” from previously reported testicular dysgenesis syndrome-like phenotype in neonatal and pre-pubertal BTP lambs. Expression of transcription factors *CREB1* (neonatal) and *BCL11A* and *FOXP2* (pre-pubertal) were significantly higher in the BTP exposed testes, with no changes seen in the adults. Increased *CREB1*, which is crucial for testes development and regulation of steroidogenic enzymes, could be an adaptive response to gestational EC exposure to facilitate the phenotypic recovery. Overall, this demonstrates that testicular effects from gestational exposure to low-level mixtures of ECs can last into adulthood, potentially impacting fertility and fecundity.

## 1. Introduction

The past 8 decades have seen a consistent decline in the reproductive health of humans and wildlife (Harrison et al., 1997). In men, this presents as reducing sperm counts and semen quality (Levine et al., 2022; Nelson and Bunge, 1974; Swan et al., 2000) concurrent with increasing rates of male reproductive disorders, including testicular cancer (Adami et al., 1994; Chia et al., 2010; Møller, 1998) and anomalies of the male external genitalia, most noticeably cryptorchidism and hypospadias (Campbell et al., 1987; Chilvers et al., 1984; Matlai and Beral, 1985; Paulozzi et al., 1999; Toppari et al., 2010). While endeavours to elucidate the underlying causes of these adverse trends in male reproductive health, now termed testicular dysgenesis syndrome (TDS) (Skakkebæk et al., 2001), have identified many contributory factors, such as malnutrition, sedentary lifestyle, and stress (Crean and Senior, 2019; Ilacqua et al., 2018), much research has focussed on the role of exposure to environmental chemicals (ECs) (Skakkebæk, 2002; Skakkebæk et al., 2022). Epidemiological evidence has shown links between gestational exposure to ECs and negative reproductive health outcomes for male offspring (Rodprasert et al., 2021), which is mirrored in animal models of gestational exposure to individual ECs, for example, phthalates (Hu et al., 2009; Repouskou et al., 2021).

While many investigative studies have concentrated on potential effects of specific chemicals or families of chemicals, there is increasing attention to the vast numbers of chemicals in the environment to which the population is constantly co-exposed. This is of concern as EC mixture effects can be seen even when the individual mixture components are administered at doses lower than their tolerable daily intake values (TDI) (Buñay et al., 2018; Kortenkamp, 2014). While efforts are being made to model more realistic EC exposure scenarios (Tsatsakis et al., 2017), it is logistically impossible in terms of both the numbers and doses of chemicals to simulate actual EC exposure by traditional component-based methodologies. The biosolids treated pasture (BTP) sheep model, however, utilises a more realistic EC exposure paradigm relative to the human populations; as biosolids (which are a commonly utilised agricultural fertiliser) are derived from domestic and industrial human waste water treatment, they contain a complex mixture of chemicals which reflects human EC exposure (Rhind et al., 2010, 2002, 2013; Venkatesan and Halden, 2014a, 2014b). Grazing of sheep on BTP during pregnancy results in measurable concentrations of many ECs in maternal and offspring organs (Bellingham et al., 2012; Filis et al., 2019; Rhind et al., 2010, 2009, 2005). A TDS-like phenotype (reduced germ cell numbers and greater rates of Sertoli-cell only (SCO) seminiferous tubules) has been previously reported in neonatal (1-day-old) and pre-pubertal (8-weeks-old) lambs whose mothers were grazed on BTP prior to and during pregnancy, with lower plasma testosterone concentrations also reported for the neonatal male offspring (Elcombe et al., 2022b, 2021). In the fetus (GD110 and GD140), across various exposure periods, fewer germ cells, Leydig cells, and Sertoli cells, and lower plasma testosterone concentrations have been reported (Lea et al., 2022; Paul et al., 2005). Results from adult rams would also suggest that BTP exposure (gestationally, and for 7 months during post-natal life, i.e., lactational / direct oral exposure) also results in a TDS-like phenotype in a subset of adult (19-months-old) rams (Bellingham et al., 2012). However, the progression of the maternal BTP exposure induced testicular phenotype seen in male offspring has not been characterised into adulthood. Testicular transcriptomic profiles have been produced from fetal, neonatal, and pre-pubertal testes, which have indicated perturbations in multiple pathways and led to evidence of alterations in transcription factor (TF) activation, specifically cAMP response element-binding protein (CREB) and hypoxia inducible factor 1 alpha (HIF1α), however, the extent of TF perturbation and the permanence of such alterations is not yet known.

The current study aimed to investigate, in parallel, the morphological and transcriptomic changes in adult (11-months-old) ram testes after pre-conceptional and gestational BTP exposure of their mothers, and to combine this data with that already collected from pre-pubertal rams from the same exposure cohort of animals, and from neonatal rams also born following maternal gestational BTP grazing. By performing analyses across these ages, we aimed to evaluate the progression of the TDS-like phenotype and persistence of TF activity over an extended period without BTP exposure.

## 2. Methods

### 2.1. Ethics statement

All procedures were carried out in line with the UK Home Office Animals (Scientific Procedures) Act (A(SP)A) 1986 regulations, under project licence PF10145DF. The project was also approved by the University of Glasgow School of Biodiversity, One Health, and Veterinary Medicine Research Ethics Committee. Animals were maintained under normal husbandry conditions at the Cochno Farm and Research Centre, University of Glasgow.

### 2.2. Experimental animals

All adult study animals were EasyCare sheep, and siblings or half siblings of the pre-pubertal rams described in Elcombe et al. (2022b), which are also used here. All neonatal animals were Aberdale sheep, and from a separate exposure cohort described in Elcombe et al., (2021). The use of animals from separate exposure cohorts was due to COVID restrictions of movement prohibiting the collection of tissues of EasyCare sheep at 1-day-old. For one month prior to mating by artificial insemination with semen from 4 rams (4 sire groupings), and for the entirety of pregnancy, ewes were maintained on either biosolids treated pasture (BTP) (biosolids exposed (B)) or pastures fertilised with inorganic fertiliser (control (C)). Pastures were fertilised twice yearly (April and September). BTPs used conventional rates of biosolids (4 tonnes / hectare) as a fertiliser, and C pastures used conventional fertiliser with equivalent amounts of nitrogen (225 kg N/ha per annum). Pregnant ewes were brought indoors two weeks prior to parturition. While maintained indoors, ewes were fed forage harvested from their respective pasture types, supplemented with concentrates as per normal husbandry practice. After birth, pre-pubertal and adult male offspring were maintained on control pastures, whereas neonatal male offspring did not leave birthing pens. At 1 day (neonatal, n = 7 control, n = 17 biosolids), 8 weeks (pre-pubertal, n = 11 control, n = 11 biosolids) or 11 months (adult, n = 11 control, n = 10 biosolids) of age, male offspring were weighed and euthanised by IV barbiturate overdose (140 mg/kg Dolethal, Vetroquinol, UK).

### 2.3. Tissue collection

At necropsy, testes were dissected. From the left testes, two slices were taken transversely from the centre, quartered, and fixed overnight in 10% neutral buffered formalin (Thermo Scientific - 16499713) before being transferred to 70% ethanol (VWR – 20821.330). Fixed sections of testes were trimmed and processed for embedding in paraffin wax for histology (Excelsior AS, Thermo Scientific). Formalin-fixed, paraffin embedded (FFPE), testicular tissues were stored at room temperature until analysis. From the right testes, transverse slices 5mm thick were taken, quartered, and frozen in liquid nitrogen prior to storage at −70°C for later RNA extraction.

### 2.4. Immuno-histochemistry

Two sections (5μm) of FFPE testicular tissues were taken for each adult animal and mounted on Polysine® coated glass slides. One section per animal underwent immuno-histochemistry (DAB staining), as previously described (Elcombe et al., 2022b), but using a rabbit anti-Sox9 antibody (Sigma-Aldrich AB5535) diluted at 1:1000 to identify Sertoli cells. The other section, and equivalent sections taken from FFPE neonatal testicular tissues, underwent fluorescent immuno-histochemistry for HIF1α as previously described (Elcombe et al., 2022b).

### 2.5. Image capture and analysis

#### 2.5.1. SOX9 immuno-histochemistry

Six images from separate areas of the lobuli testis were captured at 100x magnification (Leica DM4000B microscope, Leica DC480 digital camera) for each 11-month-old animal. Individual tubule sections entirely captured within image boundaries (n = 1483 Control, 1423 Biosolids) were manually counted. SOX9 positive Sertoli cells were identified by DAB staining and counted. Spermatogonia, spermatocytes and round spermatids were identified by circular nuclei, and elongating spermatids and spermatozoa were identified by condensed and elongated nuclei with darker haematoxylin staining. Normal seminiferous tubule sections at any stage should have a generation of elongating spermatids or spermatozoa. As per OECD guidelines (OECD, 2009), tubule sections with no or few (<5) elongating spermatids and spermatozoa were classified as showing degeneration and depletion of elongating spermatids, whereas those with many (>5) elongated spermatids and spermatozoa were classified as showing typical spermatogenesis..

#### 2.5.2. HIF1α fluorescent immuno-histochemistry

Image capture and analysis of HIF1α fluorescent immuno-histochemistry was performed as previously described (Elcombe et al., 2022b). Briefly, for each section, four images from separate areas of the lobuli testis were captured at 400x magnification (Leica DM4000B microscope, Leica DC480 digital camera). Nuclear HIF1α staining was quantified on areas outside the seminiferous tubules using the JACoP plugin (Bolte and Cordelières, 2006) for ImageJ.

### 2.6. RNA extraction, cDNA library preparation, sequencing, and data analysis

Nanopore transcriptome sequencing and analysis was performed on testicular tissues as previously described (Elcombe et al., 2022b, 2021). Differentially expressed gene (DEG) lists were subjected to gene ontology (GO) and KEGG pathway analyses in DAVID (version 6.8). The DEG list from the adult ram testes, and those previously generated from neonatal (Elcombe et al., 2021) and pre-pubertal (Elcombe et al., 2022b) ram testes, were submitted to ChEA3 for transcription factor (TF) enrichment analysis (Keenan et al., 2019), which produces a list of TFs whose gene products may be over-represented within DEG lists. Sequencing data for gene products of identified transcription factors were extracted from each dataset, geometric means of fold change values calculated, and results filtered for TFs where log2 of the geometric mean fold change was ≥1 or ≤-1 in any age group.

### 2.7. Quantitative qPCR

RNeasy Mini Kits (Qiagen – 74104) were used to extract RNA from approximately 30mg of frozen neonatal, pre-pubertal, and adult testicular tissues. A QuantiTect Reverse Transcription Kit (Qiagen – 205311) was used to degrade genomic DNA and synthesise cDNA. qPCR was performed on a Stratagene 3000 qPCR system using Brilliant II SYBR Master Mix (Agilent – 600828). Primer sequences are in Supplementary Data 1. Primer efficiencies and Ct values for each sample were calculated by regression of raw fluorescent data by PCR Miner (Zhao and Fernald, 2005). These were used in ΔΔCt analysis to calculate log2 fold change values.

### 2.8. Statistical analysis

R (version 4.1.1) base functionality was used for all calculations and statistical analyses. Data were fitted to generalised linear models using a gamma distribution and groups compared by Wald tests. Sire (adult and prepubertal cohorts) was incorporated into models to account for the genetic structure of data. The R package ggplot2 (version 3.3.5) was used to produce plots. Data reported as mean ± SD.

## 3. Results

### 3.1. Greater proportions of seminiferous tubules with degeneration and depletion of spermatids in testes of adult rams gestationally exposed to BTP

In the adults, examination of histological images (Figure 1A) revealed no differences between B and C ram testes in terms of germ cell numbers or germ cell : Sertoli cell ratios, and no incidences of SCO tubules. The testes of adult B rams contained a significantly (p = 0.0225) higher proportion of seminiferous tubule sections showing degeneration and depletion of spermatids (31.62 ± 9.92 %) compared to adult C rams (21.96 ± 6.48 %) (Figure 1B).

**Figure 1.**
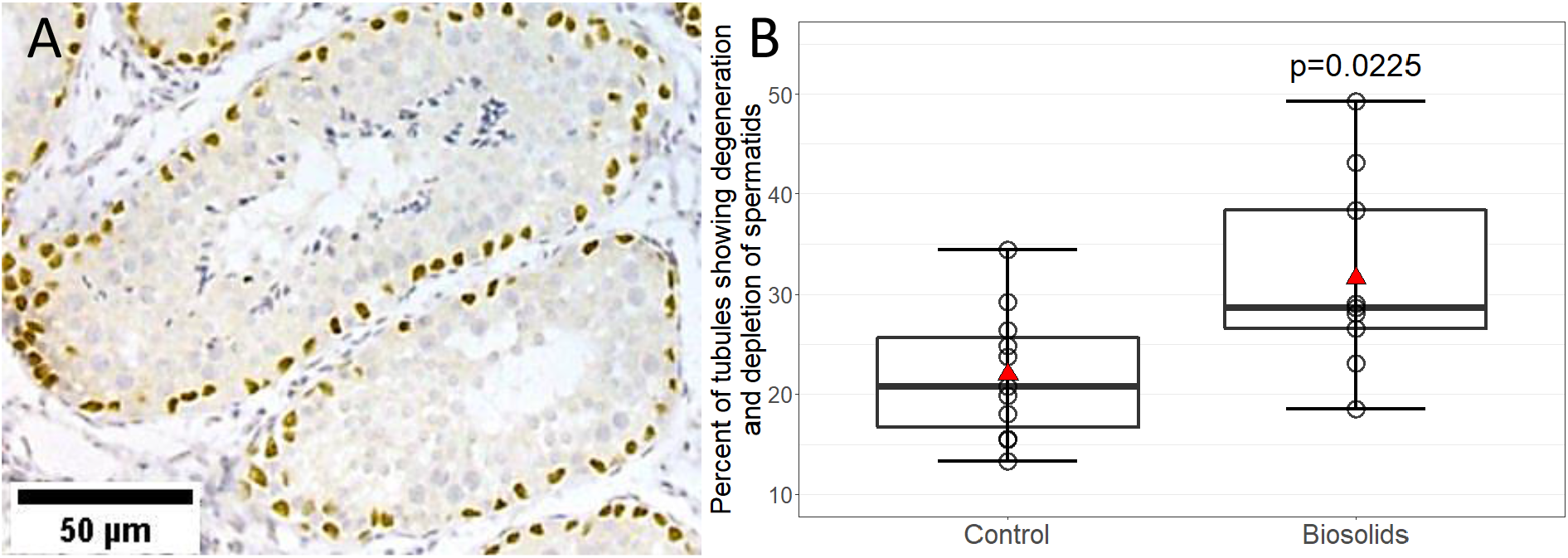
(A) Examples of seminiferous tubule sections stained with anti-SOX9 DAB and haematoxylin, showing typical morphology (top) and degeneration and depletion of elongating spermatids (bottom). (B) Biosolids rams have a higher proportion of seminiferous tubules showing degeneration and depletion of elongating spermatids. Boxes represent 25th to 75th percentile, horizontal bar indicates 50th percentile, whiskers indicate range, circles show individual data points, and red triangles show means.

### 3.2. Gestational BTP exposure alters testicular transcriptome in adult rams

Analysis of Nanopore sequenced testicular transcriptomes in the adult rams identified 1183 differentially expressed genes (DEGs) between B and C rams (562 with greater levels of expression and 621 with lower levels of expression in B relative to C). Thirty-three DEGs had a false discovery rate ≤ 0.1 (13 with greater levels of expression and 20 with lower levels of expression in B relative to C). Gene ontology (GO) analysis of the 33 DEGs indicated 5 GO terms and 1 KEGG pathway as enriched (p < 0.05) (**Error! Reference source not found.**).

**Table 1.**
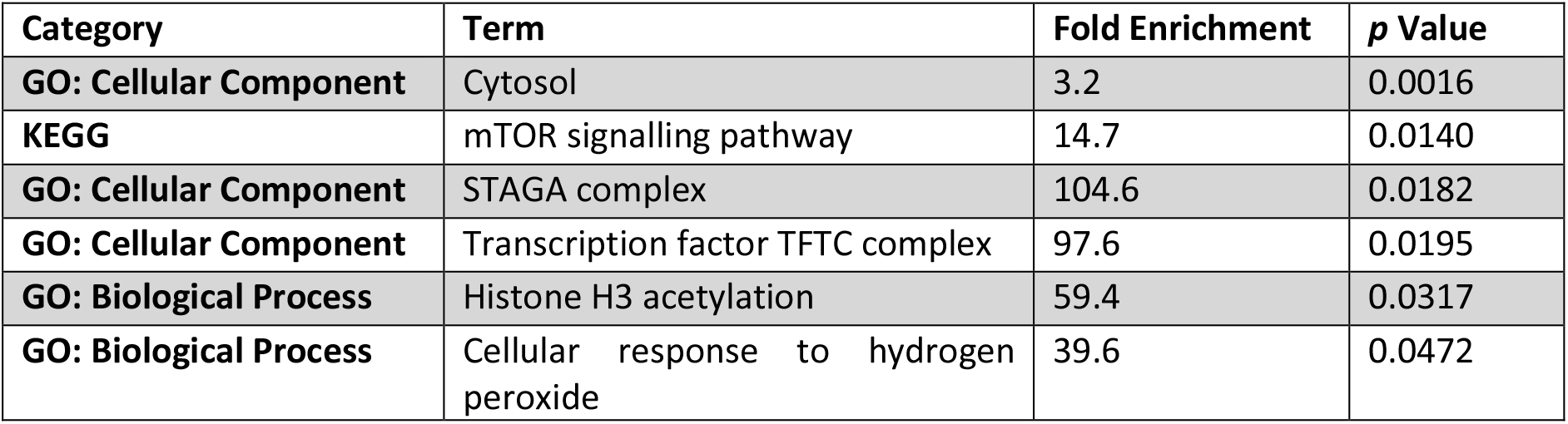
GO terms and KEGG pathways identified as potentially enriched by DAVID.

### 3.3. Extended period without exposure to BTP associated with recovery of TDS-like phenotype, deactivation of HIF1a, and the normalisation of gene expression

Testicular cell counts, proportions of SCO-tubules observed, transcriptomic findings, and HIF1α localisation data are presented in Figure 2 for the prepubertal and adult animals, which are from the same BTP exposure (immediately prior to and throughout gestation) cohort, and the neonatal lambs from a separate cohort, but which received a similar exposure (gestational). This represents a synthesis of previously generated and published data (neonatal: cell counts, SCO-tubules, and transcriptomic findings (Elcombe et al., 2021), and pre-pubertal: cell counts, SCO-tubules, transcriptomic findings, and HIF1α localisation (Elcombe et al., 2022b)), new adult data (all), and new neonatal data (HIF1α localisation). As seen in Figure 2A and B, reductions in germ cell numbers relative to Sertoli cells and the presence of SCO tubules was most pronounced in neonatal B lambs. Although from a different exposure cohort of animals, these effects of exposure were still present, although to a slightly lesser extent, in the 8-week-old B lambs. There were no differences, however, in the number of SCO tubules or the germ cell: Sertoli cell ratio in the adult rams that were derived from the same exposure cohort of animals as the 8-week-old rams.

**Figure 2.**
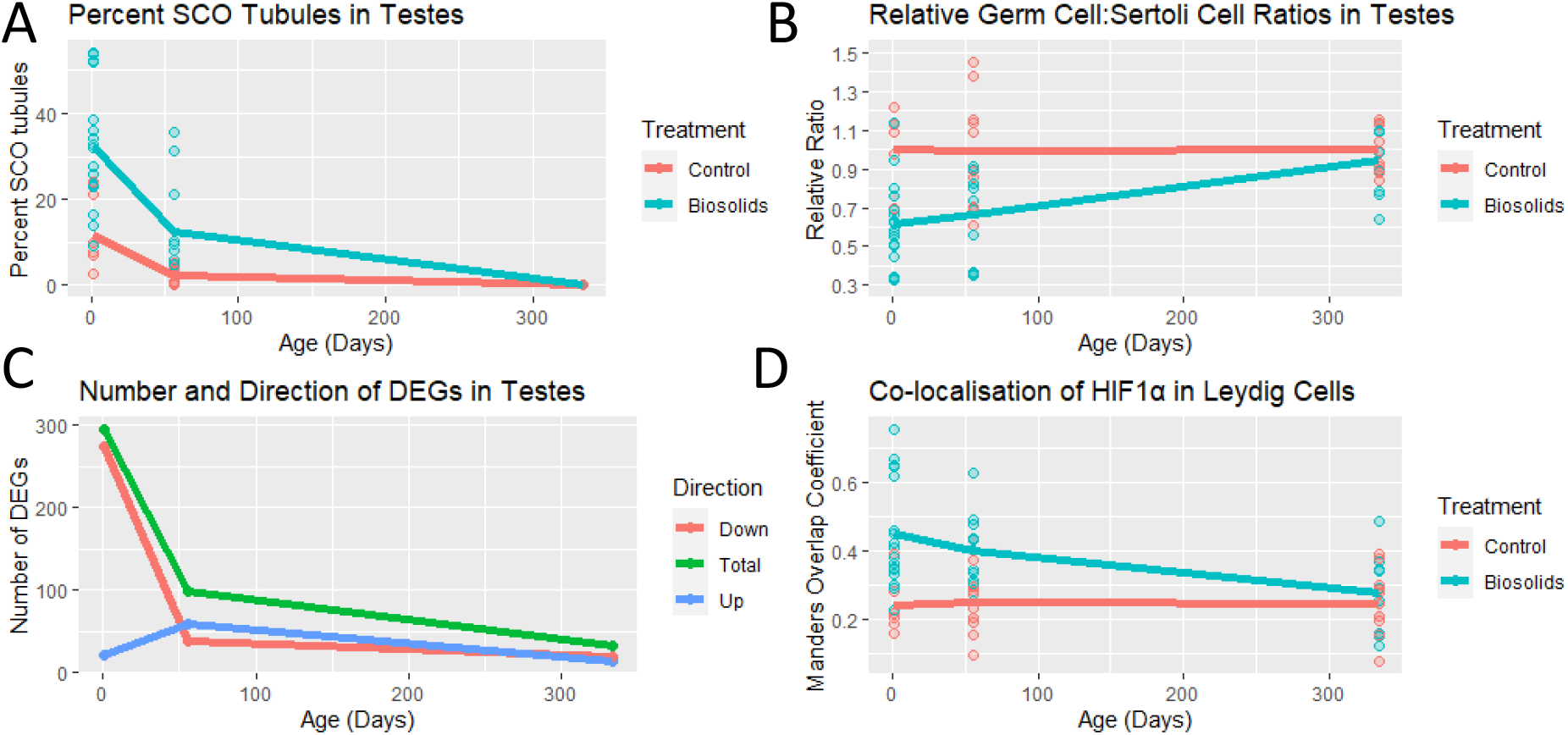
Combined data for neonatal, pre-pubertal, and adult ram testes. (A) Frequencies of SCO seminiferous tubules decrease with age. (B) Relative germ cell to Sertoli cell ratios in seminiferous tubules of biosolids rams increases towards control levels with age. (C) Numbers of genes differentially expressed between biosolids and control rams decreases with age. (D) HIF1α nuclear localisation in Leydig cells of biosolids ram testes decreases with age, returning to control levels by adulthood.

Figure 2C shows the number and direction of differentially expressed genes between the testes of B and C rams at 1-day-old (neonatal), 8-weeks-old (pre-pubertal), and 11-months-old (adult). The greatest number of DEGs observed was in the neonatal testes (296 DEGs: expression in B rams were higher for 21 genes, and lower for 275 genes, compared to C). The number of DEGs in pre-pubertal ram testes was about one third of the neonatal (99 DEGs: expression in B rams were higher for 60 genes, and lower for 39 genes, compared to C). The number of DEGs identified in the adult ram testes was one third (33 DEGs: expression in B rams were higher for 13 genes, and lower for 20 genes, compared to C) of that seen in the pre-pubertal rams, which were from the same exposure cohort and therefore only differed in age and time since exposure. There was very little overlap between DEGs identified between age groups: 7 shared between the neonatal and pre-pubertal rams, 3 shared between neonatal and adult rams, with only 1 shared between the pre-pubertal and adult rams which were derived from the same exposure cohort of animals (Supplementary data 2).

Nuclear localisation of HIF1α within Leydig cells was investigated in the neonatal maternally B exposed rams and the adult rams born to mothers exposed to BTP immediately before and throughout gestation. The data obtained was compared to the previously reported data from pre-pubertal ram testes (same exposure cohort at the adults) where activation was previously observed (Elcombe et al., 2022b). The proportion of HIF1α nuclear localisation in the C ram testes was consistent (0.241 ± 0.087, 0.251 ± 0.097, 0.249 ± 0.093) across the neonatal, pre-pubertal, and adult rams, respectively. Within the B exposed rams there was a significantly greater proportion of HIF1α nuclear localisation in the Leydig cells of the testes from the neonatal rams (0.452 ± 0.156, p = 0.0048) relative to C. This pattern was also seen in the pre-pubertal rams (0.403 ± 0.106, p = 0.0032, previously published (Elcombe et al., (2022b)), however, the adults showed no statistically significant difference (0.277 ± 0.113) in the proportion of HIF1α nuclear localisation in the Leydig cells.

To assess the impact greater or equal HIF1α nuclear localisation had on the expression of HIF1α gene products, as before in the pre-pubertal rams, qPCR was performed for *HK1, PDPK, and VEGFA* on cDNA synthesised from neonatal and adult ram testes and combined with previously published expression data for these genes in the pre-pubertal ram offspring (Elcombe et al., 2022b). Significantly greater expression of *HK1* (p = 8.4e-07) and *VEGFA* (p = 0.001) was seen in the neonatal B ram testes compared to C, and no statistically significant differences in gene expression were seen in the adult testes (Figure 3). The fold change in expression levels of *HK1* and *VEGFA* for neonatal B rams compared to same age C rams were greater than those for the pre-pubertal B ram testes (previously published in Elcombe et al., (2022b)), but expression levels in the neonatal B ram testes were of similar magnitude to the pre-pubertal B ram testes. However, curiously, *PDPK* expression was not different between B and C rams in the neonatal ram testes.

**Figure 3.**
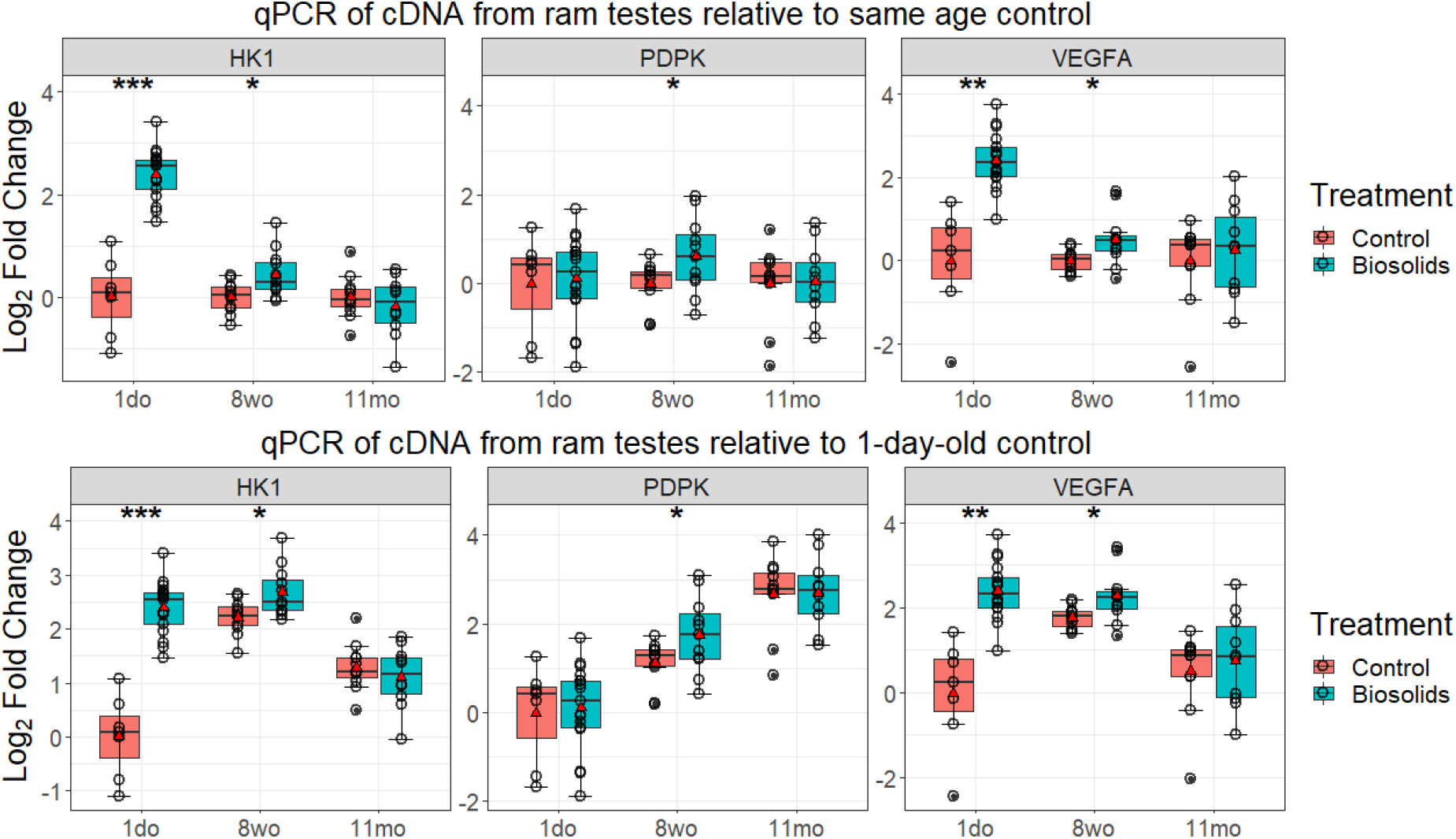
qPCR for the genes HK1, PDPK, and VEGFA in neonatal, pre-pubertal, and adult ram testes expressed as Log_2_ Fold Change in gene expression relative to same age control (top) or neonatal controls (bottom). Boxes represent 25th to 75th percentile, horizontal bar indicates 50th percentile, whiskers indicate range excluding outliers, solid filled circles show outliers, open circles show individual data points, and red triangles show means.

### 3.4. Gestational BTP exposure alters transcription factor expression in testes of neonatal and pre-pubertal lambs, but not adult rams

Transcription factor (TF) analysis of DEG lists from neonatal, pre-pubertal, and adult ram offspring by ChEA3, and subsequent interrogation of sequencing data, identified 50 TFs potentially affected by exposure (Supplementary Data 3). Of these, 43 showed lower expression of gene products in the neonatal B lamb compared to same age C, with no differences seen in the pre-pubertal or adult offspring that were derived from the same exposure cohort. 1 TF (CREB1) showed higher gene product expression in the neonatal B testes compared to same age C, and 6 TFs (BCL11A, FOSL1, FOXA1, FOXP2, GATA3, and JUND) showed altered gene product expression in either B pre-pubertal or B adult ram testes, which were of the same exposure cohort, compared to same age C (Figure 4). Of these, BCL11A, Fox1A and FOXP2 showed lower gene product expression in the neonatal B rams and higher in the pre-pubertal B rams (different exposure cohorts) compared to same age C, and GATA3 showed higher gene product expression in the pre-pubertal and adult rams (same exposure cohort) compared to same age C. To assess if differences in gene product expression were due to changes in TF expression, qPCR was performed on these TFs (Figure 5). There were no statistically significant differences in expression levels of *FOSL1, FOXA1, GATA3*, or *JUND* between B and same age C in any age group. In the neonatal testes, *CREB1* expression was significantly (p = 1.3e-04) greater in B lambs than in C. In the pre-pubertal testes, significantly greater expression of *BCL11A* (p = 0.018) and *FOXP2* (p = 0.0069) was seen in the testes of B lambs than same age C.

**Figure 4.**
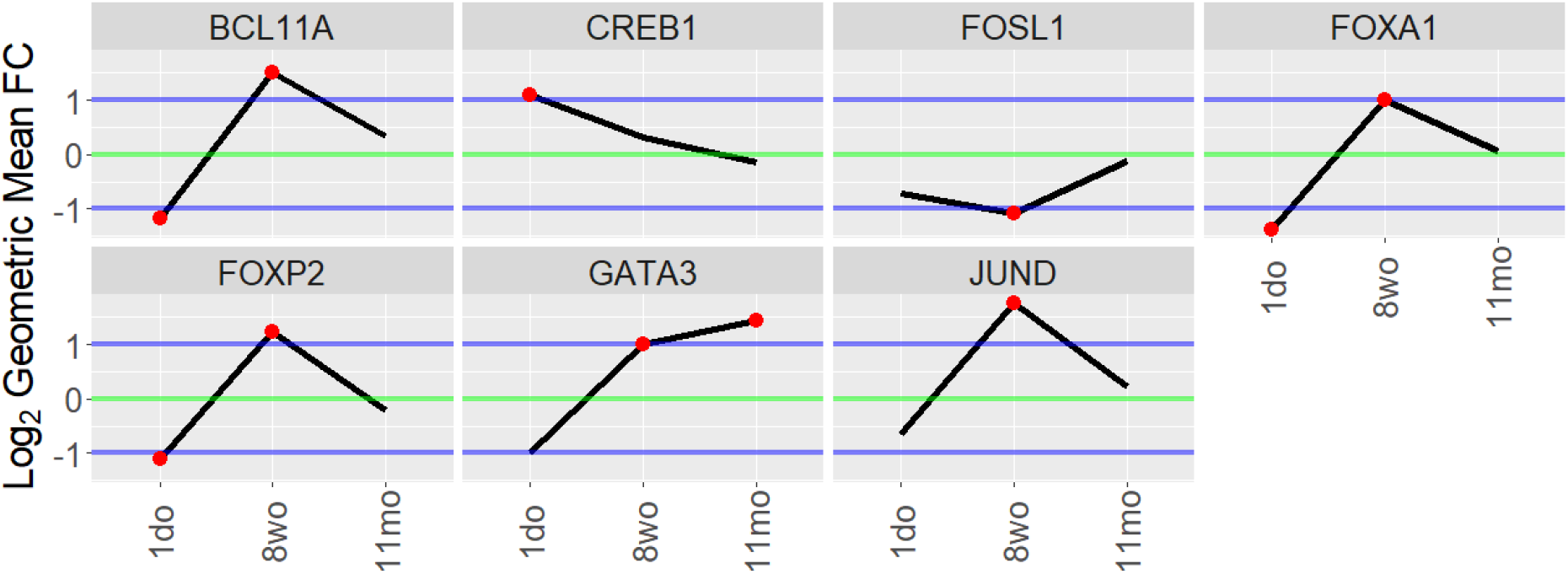
Changes in expression of gene products for transcription factors identified by ChEA3. Log_2_ of geometric means of fold change relative to same age control based on transcriptomic sequencing data, using a threshold of ±1. Red dots indicate data points which passed the threshold for inclusion.

**Figure 5.**
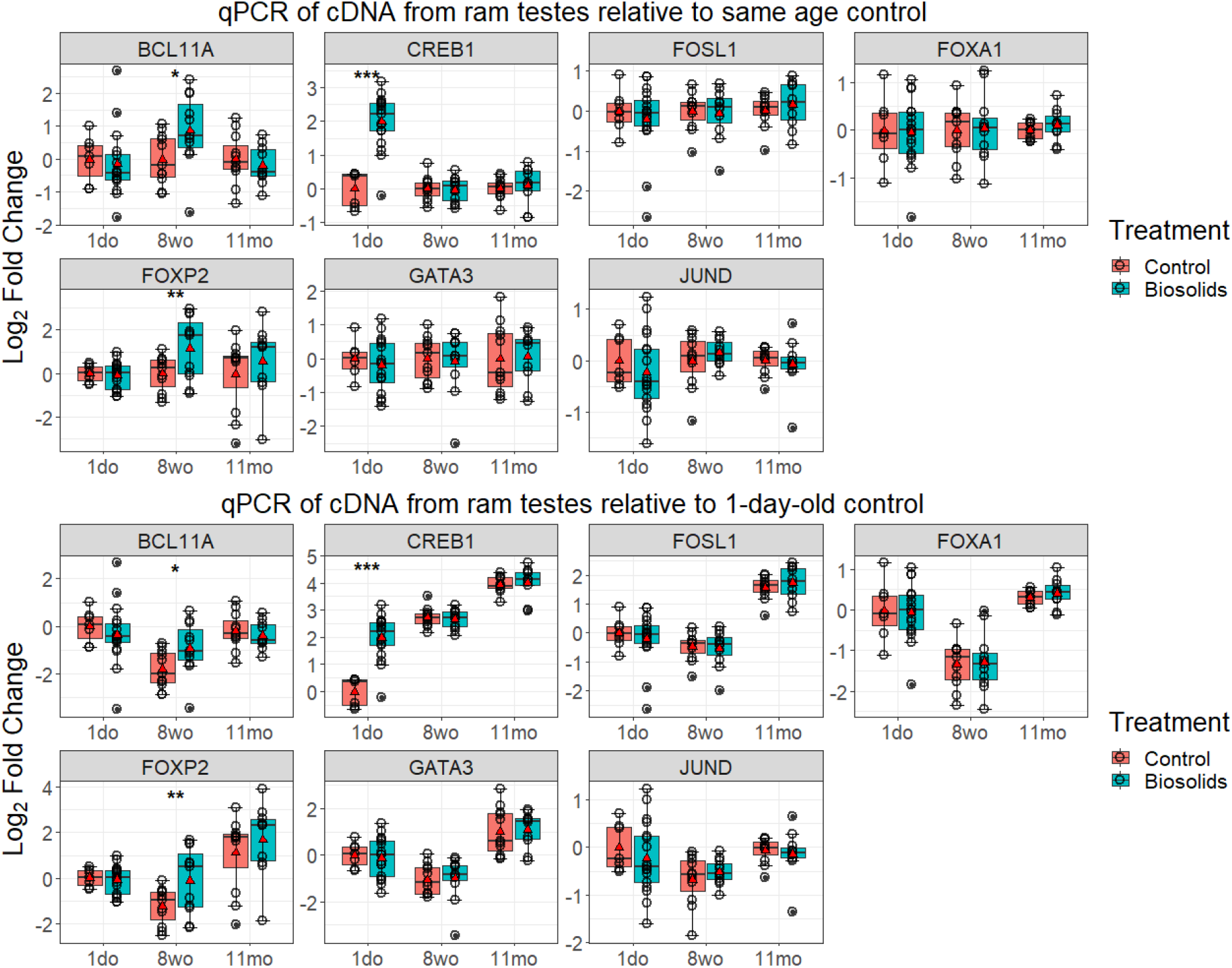
qPCR for the genes BCL11A, CREB1, FOSL1, FOXA1, FOXP2, GATA3, and JUND in neonatal, pre-pubertal, and adult ram testes expressed as Log_2_ Fold Change in gene expression relative to same age control (top) or neonatal controls (bottom). Boxes represent 25th to 75th percentile, horizontal bar indicates 50th percentile, whiskers indicate range excluding outliers, solid filled circles show outliers, open circles show individual data points, and red triangles show means.

## 4. Discussion

The long-term consequences of exposure to mixtures of ECs on male reproductive health is of continuing concern. Increasing attention towards the effects of low-dose chemical mixtures has shown that effects can often be seen following exposure to a mixture of chemicals at doses which, individually, would not be expected to elicit an effect (Elcombe et al., 2022a). Low dose gestational exposure to even simple mixtures (containing less than 10 chemicals) have been shown to cause genital malformations, alterations in testicular morphology, inhibited steroidogenesis, and impaired spermatogenesis, in male rodents (Buñay et al., 2018; Christiansen et al., 2009; Jacobsen et al., 2012). However, such component-based studies fall short of simulating the exposure scenario to which humans are constantly exposed, i.e., low concentrations of many hundreds of chemicals. The ability for gestational exposure to a complex, low-level, real-life chemical mixture to affect testicular development, with effects lasting into adulthood, is demonstrated in the present study.

Offspring of sheep grazed on BTP before and during pregnancy have shown a TDS-like phenotype (reduced germ cell numbers and greater rates of Sertoli-cell only seminiferous tubules) in neonatal rams (Elcombe et al., 2021), and in the pre-pubescent half-siblings of the adult rams presented here (Elcombe et al., 2022b). Following similar exposure, reductions in germ cell, Sertoli cell, and Leydig cell numbers has been seen in the mid-gestation (GD110) fetus (Paul et al., 2005), and in the late-gestation fetus (GD140) various exposure timings led to reductions in testes weight, total Sertoli cell numbers, and plasma testosterone concentrations (Lea et al., 2022). In the adult male offspring, there were no differences in terms of total germ cell populations and SCO tubules, which could suggest (partial) recovery from a TDS-like phenotype. Despite this apparent phenotypic recovery, adult B rams exhibited an increased proportion of seminiferous tubules showing degeneration and depletion of elongating spermatids. This phenotype is regarded as the end stage lesion of low intratesticular testosterone (OECD, 2009), which agrees with previously published observations of lower plasma testosterone concentrations in the late gestation fetus and neonatal male offspring (Elcombe et al., 2022b; Lea et al., 2022), and may have an impact on sperm production, semen quality, fertility, and fecundity. When the adult phenotype is compared to younger male offspring (neonatal and pre-pubertal) with very similar gestational BTP exposure, a lessening of morphological changes with increasing time without exposure was mirrored in both the decrease in the numbers of differentially expressed genes, and the normalisation of transcription factor expression and activity. The persistence of a TDS-like phenotype in a subset of adult males that had been grazed on BTP for 7 post-natal months (Bellingham et al., 2012) contrasts with the results seen in the adults in this study. While this phenotypic difference in adult BTP sheep testes could be a factor of susceptibility differences between breeds, as those used in Bellingham et al. (2012) were Texel sheep, it also suggests that while the TDS-like phenotype may originate during fetal development, continued EC exposure post-parturition may be required to maintain the TDS-like phenotype into adulthood. Indeed, this latter scenario is more reflective of consistent real-life exposure in humans.

The results of transcriptomic analyses are another indication of the enduring effects of fetal exposure, and while fewer DEGs were identified in the present adult ram testes than in the neonatal and pre-pubertal analyses, GO and pathway analyses still identified several changes compared to control animals. Of specific note was the identification of the mTOR signalling pathway as a site of disruption. mTOR has previously been identified as affected by EC exposure in transcriptomic analyses of fetal, neonatal, and pre-pubertal biosolids ram testes (Elcombe et al., 2022b, 2021; Lea et al., 2022). Alterations with mTOR signalling pathways due to low-level exposure should be of concern as mTOR is a crucial component of proper testicular development and spermatogenesis (Correia et al., 2020) and its disruption by chemical exposure (e.g., 4NP, DEHP, or BPA) has been shown to induce testicular autophagy in pubescent rodents (Duan et al., 2017; Fu et al., 2020; Quan et al., 2017). Additionally, mTOR is also crucial in spermatogonial stem cell maintenance, and induction or inhibition of mTOR activity can deplete the pool of spermatogonial stem cells (Hobbs et al., 2015; Xiong et al., 2015). Based on the results of our previous work in which investigations into disrupted mTOR signalling led to evidence of Hypoxia Inducible Factor 1 Alpha (HIF1α) activation and nuclear localisation in Leydig cells of biosolids pre-pubertal lambs (Elcombe et al., 2022b), this was examined in the current study in both neonatal lambs, and adult animals from the same exposure cohort as the pre-pubertal offspring. A greater proportion of HIF1α activation and nuclear localisation in Leydig cells in the neonatal lambs indicates disruption of this signalling pathway by ECs is likely to be of fetal origin. As the Leydig cells contain the highest amounts of HIF1α in the testes (Palladino et al., 2011) and, through binding site blocking, HIF1α activation can reduce the transcription of STAR, the rate limiting step in steroidogenesis, and lower testosterone production (Manna et al., 2016; Wang et al., 2019). In this respect, changes in HIF1α activation may have a role in the pathogenesis of the TDS-like phenotype seen in younger biosolids animals, where lower testosterone levels were observed in the GD140 fetus and neonatal lamb (Elcombe et al., 2021; Lea et al., 2022). However, the primary function of HIF1α is that of angiogenic and metabolic reprogramming in response to hypoxia (Child et al., 2021), and its activation could occur *via* changes in biochemical pathways (e.g. mTOR) (Dodd et al., 2015) or in direct response to xenobiotics (Bonello et al., 2007; Xia et al., 2009).

The persistent changes in activation of the TF HIF1α in biosolids animals at different ages led to the investigation of alterations in other TFs. By scrutinising transcriptomic data of TFs highlighted by ChEA3, fifty TFs were identified as potentially affected by EC exposure. Most (forty-four) had gene product expression levels altered only in the neonatal lambs, and of these all but one *(CREB1)* was associated with lower expression of gene products in the neonatal lamb. Therefore, expression levels of the six TFs identified in the other age groups, and *CREB1*, were quantified to investigate if the increased gene product transcription was a result of increased TF transcription. Higher transcription levels of *CREB1* in the neonatal, and *BCL11A* and *FOXP2* in the pre-pubertal testes matched the increased transcription of gene products of those TFs identified within the sequencing data for those age groups. While there is no literature on the roles of BCL11A or FOXP2 within the testes, Cyclic AMP (cAMP) signalling and CREB (cAMP-response element binding protein) have previously been identified as affected by EC exposure in the neonatal biosolids lamb (Elcombe et al., 2021), and are known to play important roles within the testes. Within Sertoli cells, cAMP signalling and CREB are crucial in testicular development and spermatogenesis (Don and Stelzer, 2002), and within Leydig cells CREB regulates the expression of important steroidogenic genes (Kumar et al., 2018). It is therefore possible that the activation and increased transcription of *CREB1* in the biosolid ram testes is an adaptive response to EC exposure.

A challenge and limitation in the interpretation of the present results are that two age groups (pre-pubertal and adult) are from the same exposure cohort, are of the same breed of sheep, and are full or half-siblings, whereas the other age group (neonatal) were different in all these respects. As the chemical contents of biosolids has batch variability, and chemical uptake varies based on factors of soil content (Clarke and Smith, 2011; Rhind et al., 2013; Zhang et al., 2015), there were undoubtedly differences in chemical composition between the exposure received by the neonatal rams and that to which the pre-pubertal and adult rams received. The genetic differences between Aberdale (neonatal rams) and EasyCare (pre-pubertal and adult rams) sheep is another variable of unknown impact on the results, which could affect susceptibility or resistance to exposure-induced effects and must be considered while interpreting. However, that the pre-pubertal rams morphologically resembled the neonatal rams more than their older exposure cohort counterparts, the adults, and that the prepubertal and neonatal rams showed very similar HIF1α activation patterns, these concerns were not considered to be major factors with regards to the present study. Therefore, this allowed us to compare the effects of exposure over multiple ages, which allows considerations of directionality and longevity, which is a strength of the present study. An additional strength comes from using a more translationally relevant animal system, rather than traditional laboratory rodents. Sheep are precocial with organ development more similar at birth to humans than altricial rodents and are more physiologically like humans in terms of reproductive cycle, gestational periods, lifespan, steroidal biosynthesis, and start and duration of puberty. Crucially, similarities in testis development between sheep and humans, in terms of hypothalamic–pituitary–gonadal axis function onset, plasma androgen and anti-Mullerian hormone levels, genital tubercle formation, and external genitalia differentiation, are much greater than for other species, including rats and mice (O’Shaughnessy and Fowler, 2011).

It is well recognised that fetal development is a period of increased risk to xenobiotic induced toxicity, especially that mediated by endocrine disruption. The present study exemplifies this, evidencing that exposure to an environmental chemical mixture, at realistically low doses, during gestation alone is sufficient to produce observable morphological and molecular effects in the testes, which persist into adulthood. Differences between the adult rams in this study and adult rams from a similar study, with a period of post-natal exposure, indicates a crucial period of life whereby, with continued exposure, TDS-like traits persist into adulthood. This also suggests adverse effects evident in early life are not permanent and may be at least partially recoverable, dependant on removal from the source of exposure. Increased HIF1α activation in Leydig cells is shown to be present from birth in exposed offspring, which may be linked to lowered steroidogenesis during fetal development and early life. Increased transcription of *CREB1* could be a compensatory mechanism against this action, and therefore may be important in the partial recovery observed. The current findings add to the increasing body of evidence suggesting that exposure to real-world levels of environmental chemical mixtures during pregnancy may be having a negative impact on the reproductive health of male offspring, contributing to declining male reproductive health, including sperm counts, semen quality, fertility and fecundity.

## Acknowledgments

We are grateful to the staff at Cochno Farm and Research Centre for their technical assistance.

## Supplementary data 1

**Supplementary Data 1.**
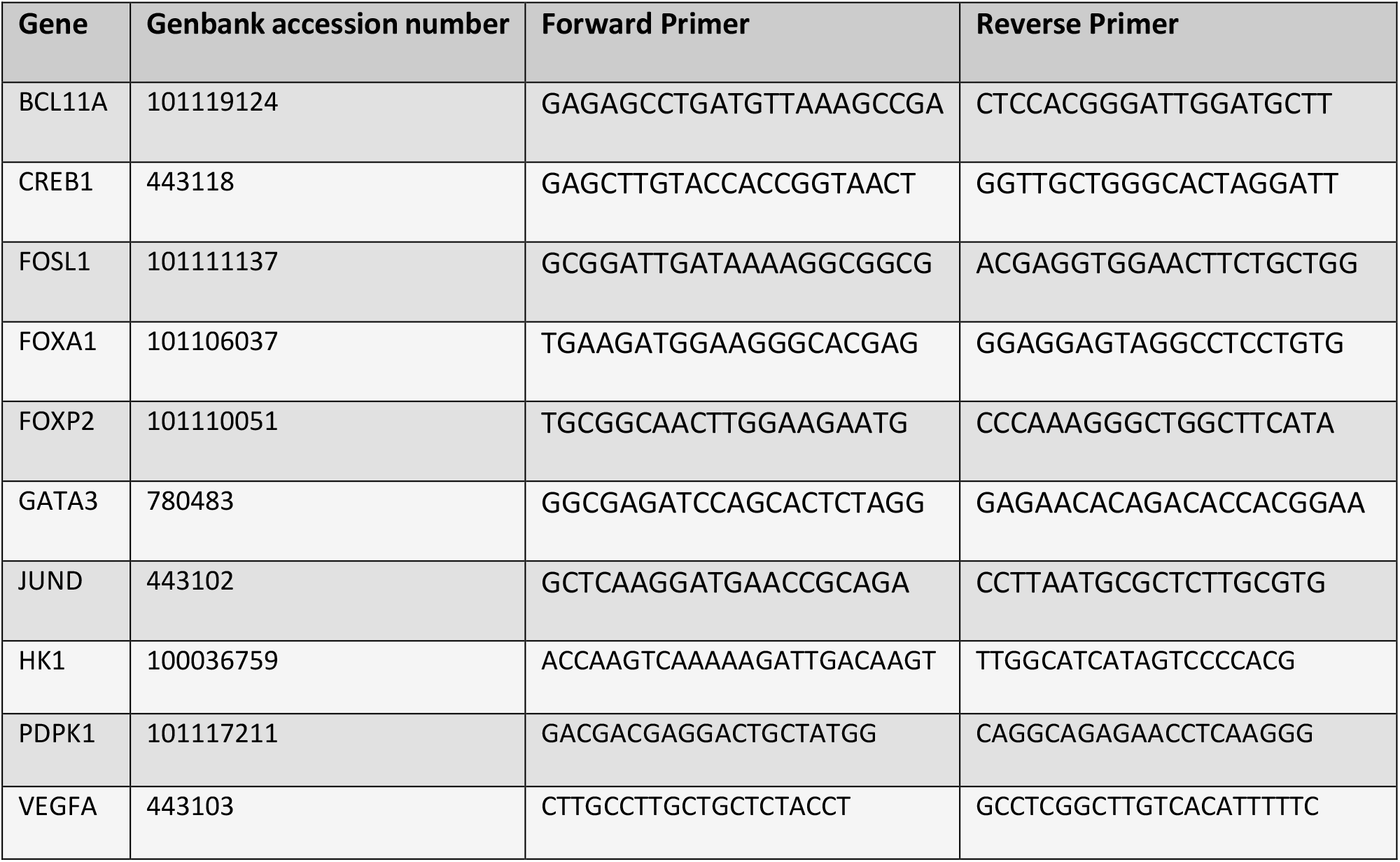
Table of qPCR primer details

## Supplementary data 2

**Supplementary Data 2.**
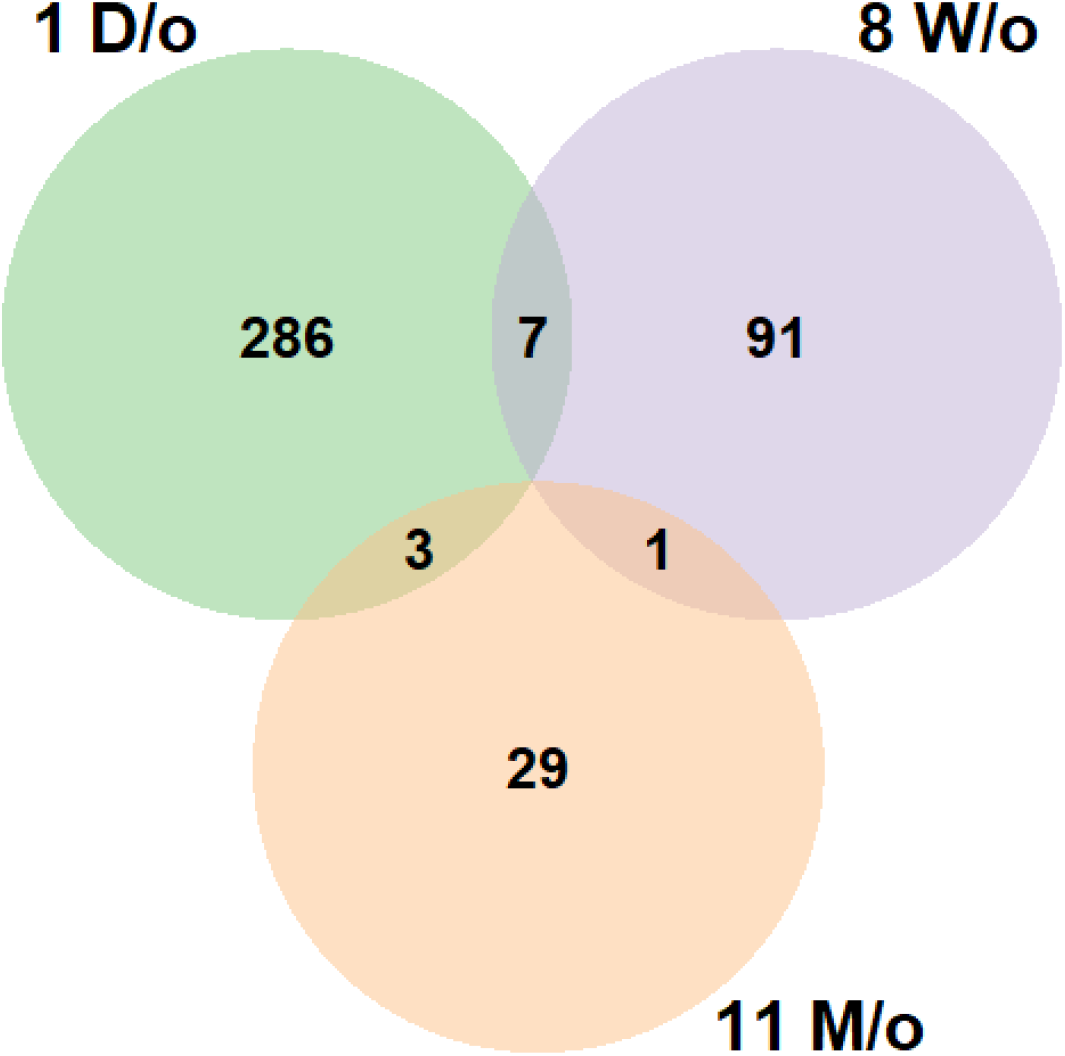
VENN diagram showing DEGs identified in neonatal (1D/o), pre-pubertal (8W/o), and adult (11M/o) ram offspring testes.

## Supplementary data 3

**Supplementary Data 3.**
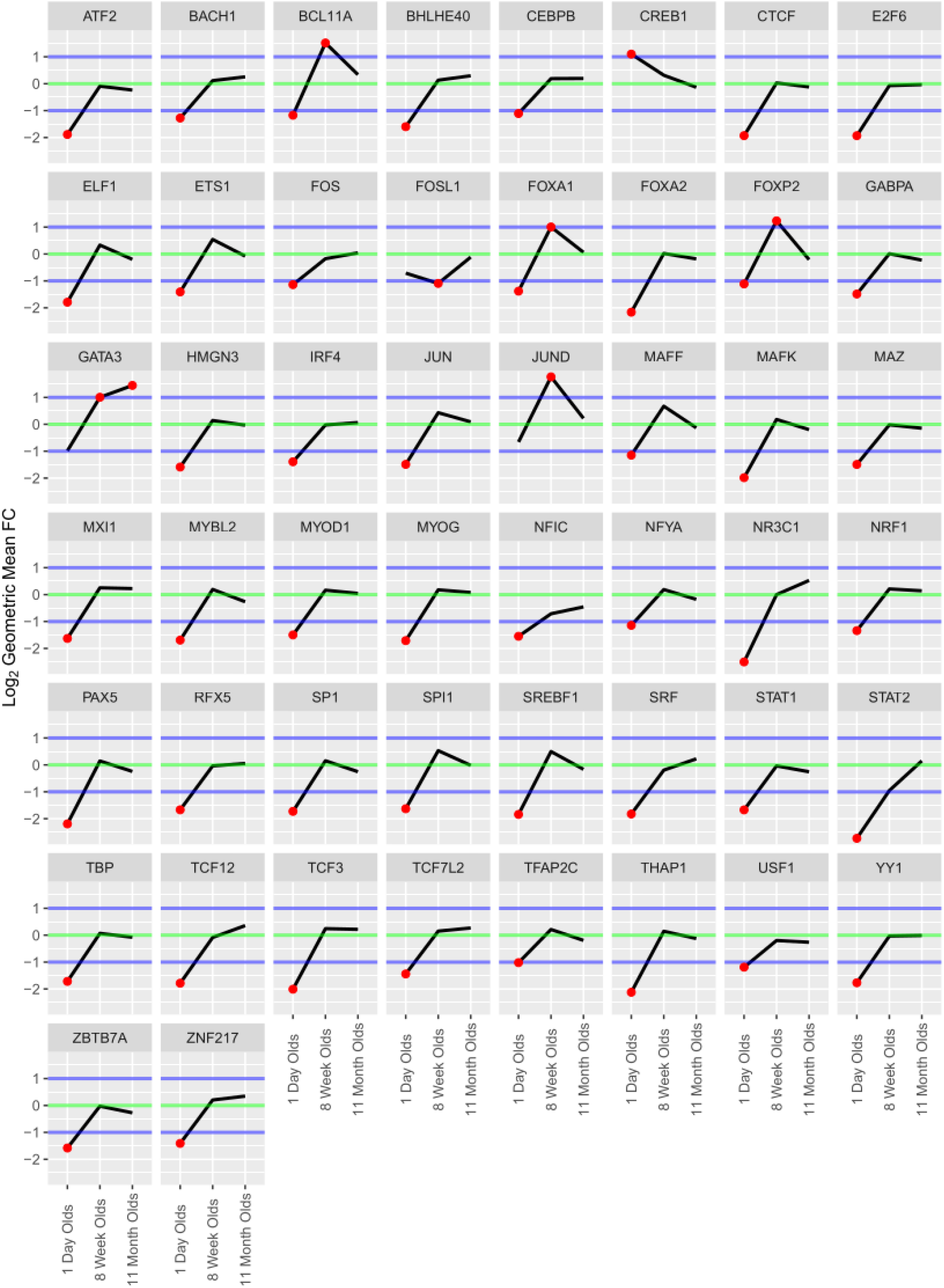
TFs which were identified as enriched by ChEA3 analysis, which had log_2_ geometric mean fold change of genes products ≥1 or ≤-1 in any age group. Red dots indicate where gene product expression levels passed this threshold.

## Abbreviations

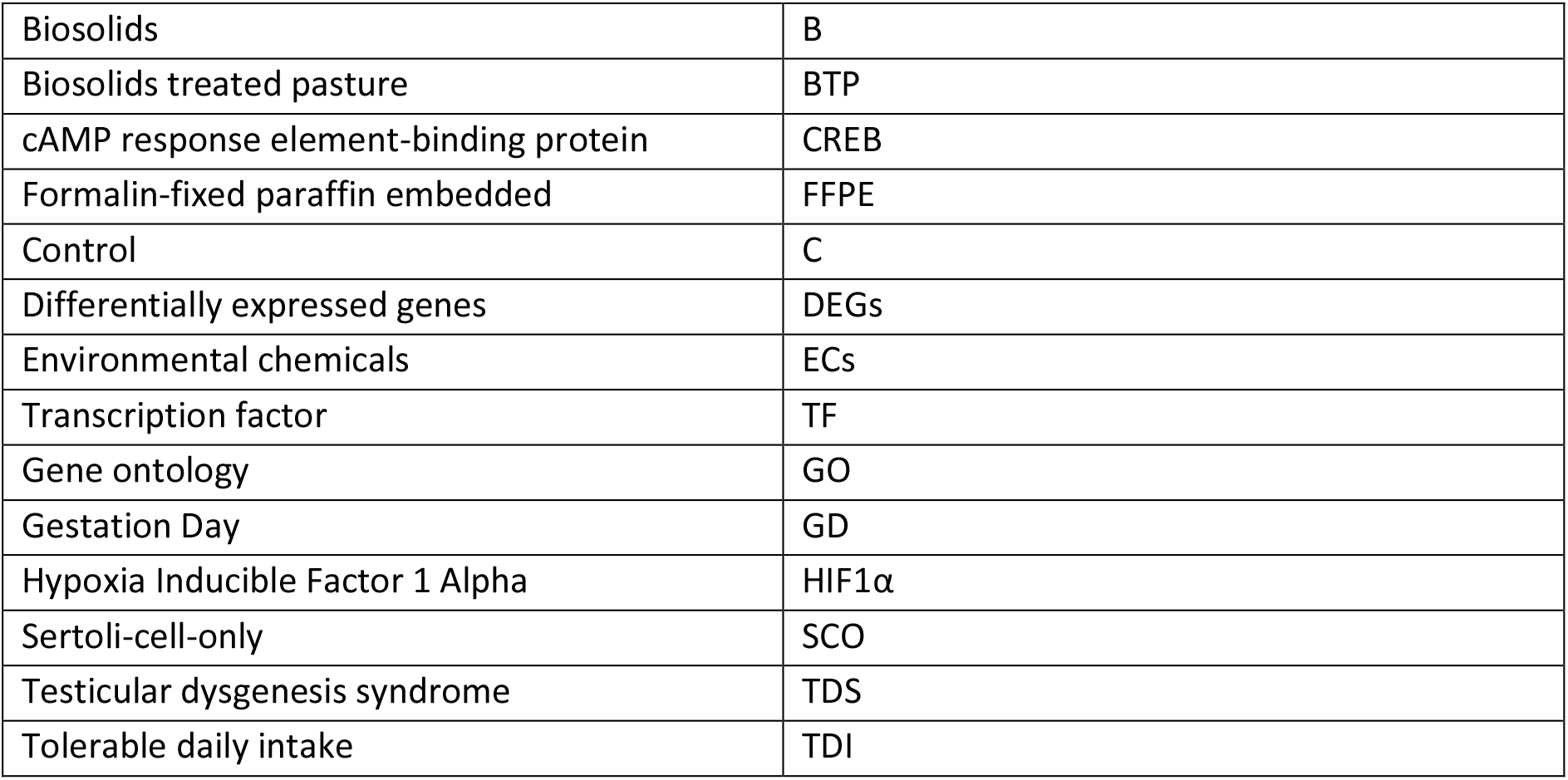

