## Supplementary data for "Developmental exposure to a real-life environmental chemical mixture alters testicular transcription factor expression in neonatal and pre-pubertal rams, with morphological changes persisting into adulthood"

Supplementary data 1

Supplementary Data 1. Table of qPCR primer details

| **Gene** | **Genbank accession number** | **Forward Primer** | **Reverse Primer** |
| --- | --- | --- | --- |
| BCL11A | 101119124 | GAGAGCCTGATGTTAAAGCCGA | CTCCACGGGATTGGATGCTT |
| CREB1 | 443118 | GAGCTTGTACCACCGGTAACT | GGTTGCTGGGCACTAGGATT |
| FOSL1 | 101111137 | GCGGATTGATAAAAGGCGGCG | ACGAGGTGGAACTTCTGCTGG |
| FOXA1 | 101106037 | TGAAGATGGAAGGGCACGAG | GGAGGAGTAGGCCTCCTGTG |
| FOXP2 | 101110051 | TGCGGCAACTTGGAAGAATG | CCCAAAGGGCTGGCTTCATA |
| GATA3 | 780483 | GGCGAGATCCAGCACTCTAGG | GAGAACACAGACACCACGGAA |
| JUND | 443102 | GCTCAAGGATGAACCGCAGA | CCTTAATGCGCTCTTGCGTG |
| HK1 | 100036759 | ACCAAGTCAAAAAGATTGACAAGT | TTGGCATCATAGTCCCCACG |
| PDPK1 | 101117211 | GACGACGAGGACTGCTATGG | CAGGCAGAGAACCTCAAGGG |
| VEGFA | 443103 | CTTGCCTTGCTGCTCTACCT | GCCTCGGCTTGTCACATTTTTC |

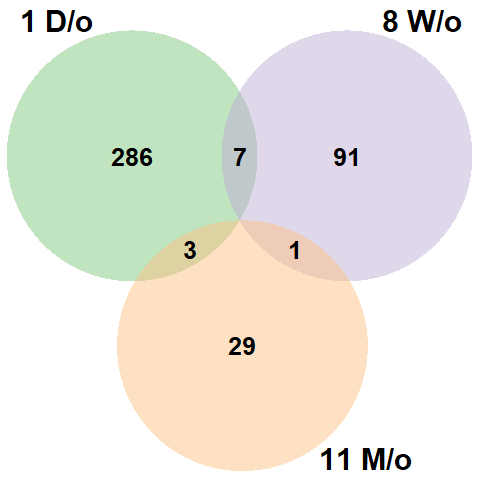
Supplementary data 2

Supplementary Data 2. VENN diagram showing DEGs identified in neonatal (1D/o), pre-pubertal (8W/o), and adult (11M/o) ram offspring testes.

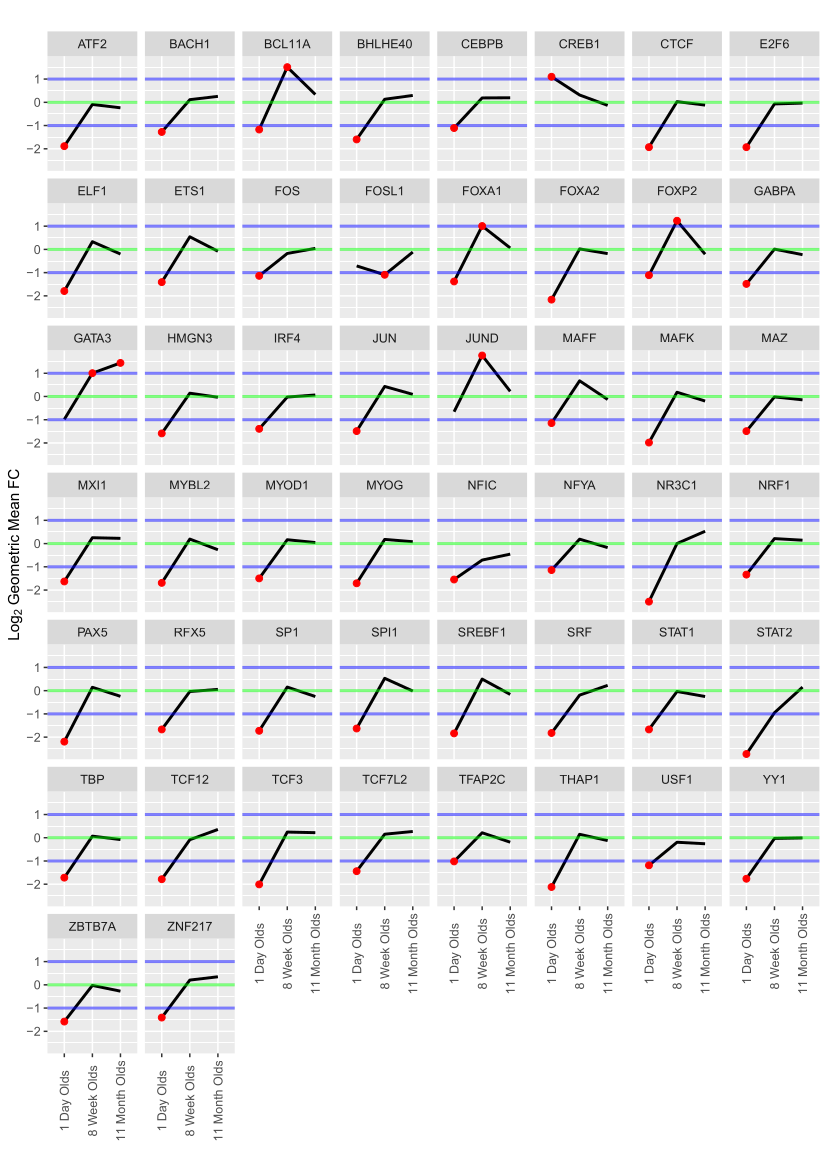
Supplementary data 3

Supplementary Data 3. TFs which were identified as enriched by ChEA3 analysis, which had log_2_ geometric mean fold change of genes products ≥1 or ≤-1 in any age group. Red dots indicate where gene product expression levels passed this threshold.
